# A neurocomputational theory of action regulation predicts motor behavior in neurotypical individuals and patients with Parkinson’s disease

**DOI:** 10.1101/2022.04.20.488864

**Authors:** S. Zhong, J. Choi, N. Hashoush, D. Babayan, M. Malekmohammadi, N. Pouratian, V. N. Christopoulos

## Abstract

Surviving in an uncertain environment requires not only the ability to select the best action, but also the flexibility to withhold inappropriate actions when the environmental conditions change. Although selecting and withholding actions have been extensively studied in both human and animals, there is still lack of consensus on the mechanism underlying these action regulation functions, and more importantly, how they inter-relate. A critical gap impeding progress is the lack of a computational theory that will integrate the mechanisms of action regulation into a unified framework. The current study aims to advance our understanding by developing a neurodynamical computational theory that models the mechanism of action regulation that involves suppressing responses, and predicts how disruption of this mechanism can lead to motor deficits in Parkinson’s disease (PD) patients. We tested the model predictions in neurotypical individuals and PD patients in three behavioral tasks that involve free action selection between two opposed directions, action selection in the presence of conflicting information and abandoning an ongoing action when a stop signal is presented. Our results and theory suggest an integrated mechanism of action regulation that affects both action initiation and inhibition. When this mechanism is disrupted, motor behavior is affected, leading to longer reaction times and higher error rates in action inhibition.

**Author Summary:** Humans can rapidly regulate actions according to updated demands of the environment. A key component of action regulation is action inhibition, the failure of which contributes to various neuropsychiatric disorders. When faced with multiple choices, dealing with conflicting information, or current actions become inappropriate or unwanted, we should be able to pause or completely abandon actions. Despite extensive efforts to understand how the brain selects, pauses, and abandons actions based on environmental demands, the mechanisms underlying these action regulation functions and, perhaps more importantly, how they inter-relate remain elusive. Part of this challenge lies in the fact that these mechanisms were rarely explored together, making it difficult to develop a unified theory that explains the computational aspects of action regulation functions. The current study introduces a large-scale model that better characterizes the computations of action regulation functions, how they are implemented within brain networks that involve frontal, motor and basal ganglia (BG) circuits, and how disruption of these circuits can lead to deficits in motor behavior seen in Parkinson’s disease (PD).The model was developed by studying the motor behavior of healthy individuals and PD patients in three motor tasks that involve action inhibition. Overall, the model explains many key aspects on how the brain regulates actions that involve inhibitory processes, opening new avenues for improving and developing therapeutic interventions for diseases that may involve these circuits.

## 1 Introduction

Surviving in an uncertain environment requires not only the ability to accurately and rapidly select the best action, but also the flexibility to abandon obsolete actions when they are rendered unwanted or inappropriate. How actions are initiated and regulated is a fundamental neurobiological question that is of high impact for understanding how the human brain functions. A key component of action regulation is inhibiting actions, which when abnormal contributes to neuropsychiatric diseases, such as Parkinson’s disease (PD), obsessive-compulsive disorder (OCD), and others [1–5]. Action inhibition occurs in at least 3 ways: (a) action selection – selecting one action requires suppressing alternative motor plans, (b) decision conflict – choosing in the presence of conflicting information requires suppressing alternative actions to buy more time to make a correct decision and (c) outright stopping – inhibiting a response when it is rendered inappropriate.

Over the past years, a number of studies attempted to characterize the mechanism of action regulation that involves action inhibition under different experimental paradigms. A recent cognitive theory suggests that action selection occurs through a competitive process between movement plans [6–10]. According to this theory, in situations affording more than one alternatives, animals prepare multiple actions in parallel that compete for selection through mutual inhibitory interactions before choosing to execute one. This affordance competition theory received empirical support from neurophysiological investigations in the sensorimotor areas of non-human primates (NHPs) showing that the brain encodes parallel reach, grasp and saccade plans before the animals select between them [11–13]. It is consistent with the continuous flow model of perception, which suggests that response preparation can begin even before the goal is fully identified and a decision is made [14–16]. In addition, psychophysical support for this theory comes from the “go-before-you-know” experiments, in which individuals had to initiate reaching or saccade movements towards multiple potential targets, without knowing the actual location of the goal [17–19]. The individuals compensate for the goal location uncertainty by aiming towards an intermediate location, a strategy consistent with an averaging of multiple competing action plans.

Recent studies have also explored the mechanisms underlying pausing or abandoning actions using functional MRI [20, 21], local field potential (LFP) recordings [22, 23], electroencephalography (EEG) recordings [24, 25], as well as single-unit recordings in humans [26, 27], non-human primates (NHPs) [28, 29] and rodents [30]. The basal ganglia (BG), and in particular the subthalamic nucleus (STN), has been functionally implicated in action regulation functions, but in association with distinct frontal areas, such as the primary motor cortex (M1), the premotor cortex (preMC), the pre-supplementary motor area (preSMA) and the right inferior frontal gyrus (rIFG) [31–34]. In a sense, STN is activated when a stop signal is detected, as well as when conflicting information is presented, to rapidly suppress ongoing or planned actions [35–37].

Despite the significant contribution of these studies on understanding how the brain selects between competing options, deals with conflicting information, and stops planned or ongoing actions during decisions, the mechanisms of these action regulation functions and their inter-relations remain elusive. Part of this challenge lies in the fact that previous studies rarely explore these functions together, making it difficult to develop a unified and integrated theory of action regulation. The current study aims to advance our understanding on the mechanism underlying action regulation and how disruption of this mechanism can lead to deficits in motor behavior exhibited in Parkinson’s disease (PD). To address these questions, we trained neurotypical individuals and PD patients to perform three motor tasks that involve motor decision between two opposed directions, action selection in the presence of conflicting information and suppression of unwanted motor responses when a stop signal is presented. To elucidate the action regulation mechanism in control and disease state, we modeled the tasks within a neurodynamical computational framework that combines dynamic field theory with stochastic optimal control theory, and simulates the processes underlying selection, planning, initiation and suppression of actions [38, 39].

Our study presents the first unified theory on action regulation that involves response inhibition, providing important predictions on how the disruption of major nodes, such as STN, can deteriorate motor performance leading to longer reaction times in motor decisions and higher error rates when stopping ongoing actions. Additionally, the neurodynamical theory provides a potential explanation on why PD patients exhibit longer reaction times than neurotypical individuals even in the lack of competing alternatives or conflicting information in motor decisions. Overall, our findings shed light on how the brain regulates actions that involve inhibitory processes, opening new avenues for improving and developing therapeutic interventions for diseases that may involve these circuits.

## 2 Results

### 2.1 Experimental paradigms

Participants were instructed to perform reaching movements using a 2-dimensional joystick under three experimental paradigms: i) decision-making task (action selection), ii) Eriksen flanker task (decision conflict) and iii) stop-signal task (outright stopping) (Fig.1). In the decision-making task, participants had to respond to arrow stimuli presented on a computer screen by freely moving the joystick towards the left or right direction. Choice trials were interleaved with instructed trials in which all arrows pointed to the same direction. In the Eriksen flanker task, flanking arrows were presented on the screen, all pointing to the same direction. A target arrow was then presented to indicate the direction to move, either in the same (no conflict, congruent trials) or opposite (conflict, incongruent trials) direction as the flanking arrows. Finally, in the stop-signal task, the participants were instructed to reach towards the direction of the arrows. In a minority of trials, the color of the arrows turned red after a short delay, and the action had to be abandoned immediately.

**Fig. 1.**
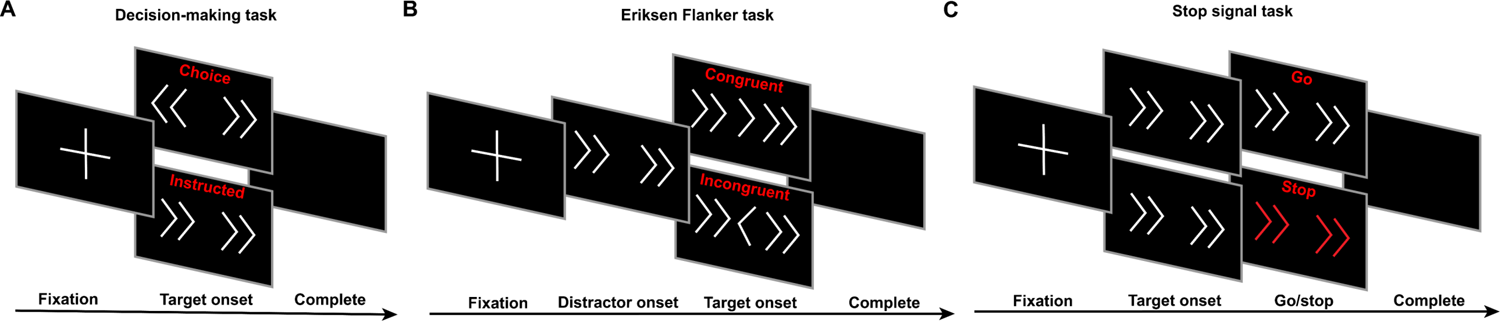
Experimental setup for action regulation tasks that require action inhibition. (A) Decision-making task, including instructed and choice trials. (B) An arrow version of the Eriksen Flanker task, including congruent (the flanker arrows point to the same direction as the central arrow) and incongruent (the flanker arrows point to the opposite direction from the central arrow) trials. (C) A stop-signal task with instructed trials. Individuals are prompted to stop the action when the arrows turn red.

### 2.2 Motor behavior of neurotypical individuals and PD patients in action regulation tasks

We computed the reaction time (RT) for initiating an action as the time interval between the presentation of the target arrows on the screen and the initiation of the reaching movement. We found that in the decision-making task, choice trials had longer RT than instructed trials in both populations (Fig. 2A) (p<0.001, two-way ANOVA). Interestingly, although the neurotypical participants responded faster than the PD patients in the instructed trials (p<0.001, two-way ANOVA), we found no significant difference in RT between the two groups in the choice trials (p=0.878, two-way ANOVA), (Fig.2A). In the Eriksen flanker task, both groups exhibited shorter RT in the congruent trials than in the incongruent trials (Fig. 2B) (p<0.001 for both neurotypical participants and PD patients, two-way ANOVA). However, PD patients had slower responses than neurotypical participants in both congruent and incongruent trials (p<0.01 for congruent trials, p<0.05 for incongruent trials, two-way ANOVA). Regarding the stop-signal task, interestingly, we found that neurotypical participants had slower responses than PD patients in the go trials (Fig.2C) (p<0.001, two sample t-test). In particular, the neurotypical group seems to have strategically slowed down their responses in the go trials by 233 ms on average in order to be more successful in inhibiting their response in stop trials (p<0.001, two sample t-test on RT between instructed trials and go trials for the neurotypical population). On the other hand, PD patients exhibited much subtler modification of their response between instructed trials (decision-making task) and go trials (stop-signal task) - the reaction time for go trials increased only by 47 ms on average compared to instructed trials (p<0.001, two sample t-test), suggesting that the anticipation of the stop signal had smaller effect on their motor planning behavior.

**Fig. 2.**
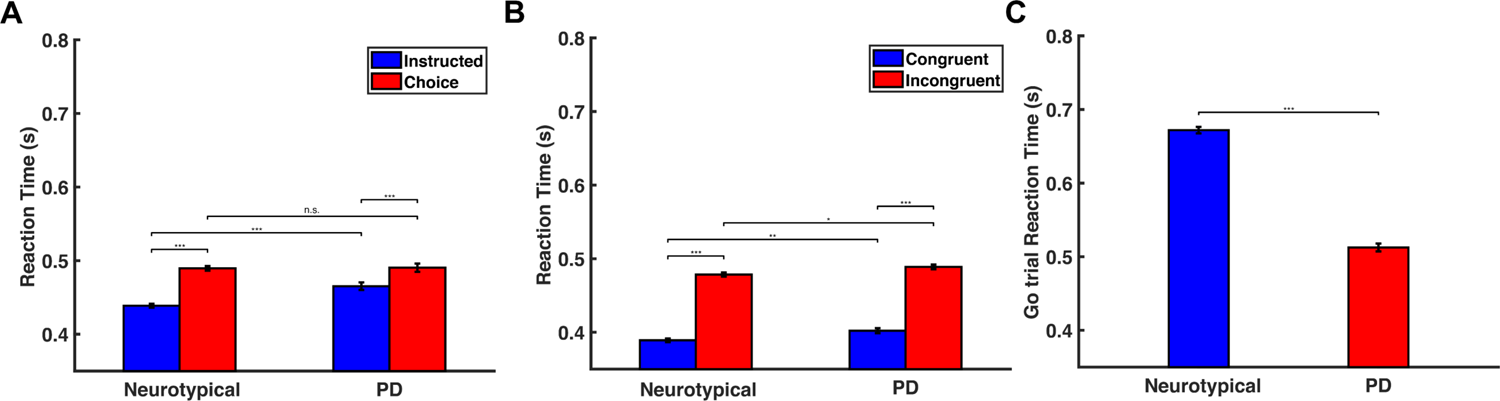
Behavioral findings from the decision-making task, the Eriksen flanker task and the stop-signal task. (A) Bar plots of the RT for neurotypical individuals (Neurotypical) and PD patients (PD) in the instructed and choice trials of the decision-making task. Error bars correspond to standard error (SE). (B) Bar plots of the RT for neurotypical individuals (Neurotypical) and PD patients (PD) in the congruent and incongruent trials of the Eriksen flanker task. Error bars correspond to standard error (SE). (C) Bar plots of the RT for neurotypical individuals (Neurotypical) and PD patients (PD) in the go trials of the stop-signal task. Error bars correspond to standard error (SE).

These findings predict that PD patients will perform worse in stop trials than neurotypical participants, since a lower probability of stopping has often been associated with faster responses in go trials [40–42]. To test this hypothesis, we computed the probability to stop an action for different stop-signal delay (SSD) values across all participants in each group. The results showed that the probability to successfully stop an action was inversely correlated with SSD, and consistent with the hypothesis, PD patients exhibited lower probability of stopping an action compared to neurotypical individuals (Fig.3).

**Fig. 3.**
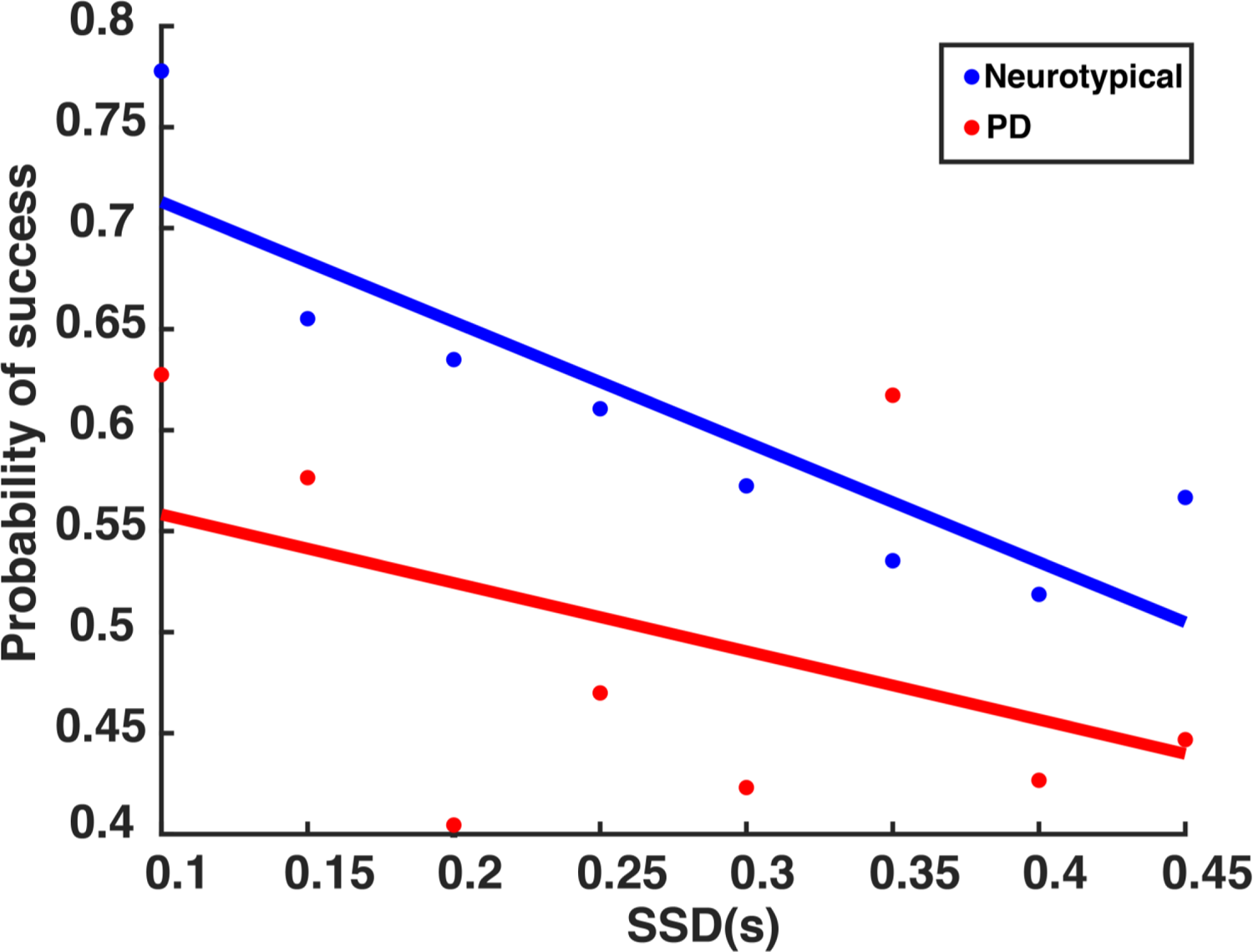
Probability to successfully stop an action as a function of the stop signal delay (SSD). The probability to successfully stop an action as a function of the SSD for neurotypical individuals (Neurotypical, blue) and PD patients (PD, red).

### 2.3 An integrated neurodynamical theory of action regulation predicts motor behavior

Our findings require a computational theory that could explain the mechanism of action regulation that involves inhibition and predicts how disruption of this mechanism can lead to motor impairments in PD patients. Building on our previous successful work in modeling visuomotor tasks [38, 39], we developed a neurodynamical theory to unify the action regulation mechanism that involves inhibition. The theory builds on the affordance competition hypothesis, according to which multiple actions are formed concurrently and compete over time until one has sufficient evidence to win the competition [6, 7, 12]. It combines dynamic neural field (DNF) theory [43, 44] with stochastic optimal control theory [45, 46] and its architectural organization is illustrated in Fig.4. Each DNF field simulates the dynamic evolution of firing rate activity of a network of neurons over a continuous space with local excitation and surround inhibition. It consists of 181 neurons - with exception of the context signal field and the pause field - and each of them has a preferred direction between 0*^◦^* and 180*^◦^*. The “spatial sensory input” field encodes the angular representation of the competing actions (i.e., left vs. right movements in our study). The “expected outcome” field encodes the expected reward for reaching to a particular direction. The outputs of these two fields send excitatory projections (green arrows) to the “reach planning” field in a topological manner. The “reach cost” field encodes the effort cost required to implement an action at a given time and state. The reach cost field sends inhibitory projections (red arrow) to the reach planning field to penalize high-effort actions. For instance, an action that requires changing of moving direction is more “costly” than an action of keeping going in the same direction. Although the cost field does not have a critical role in this study, since all planning actions are associated with about same effort, it is required for generating reaching movements from the optimal control part of the model.

**Fig. 4.**
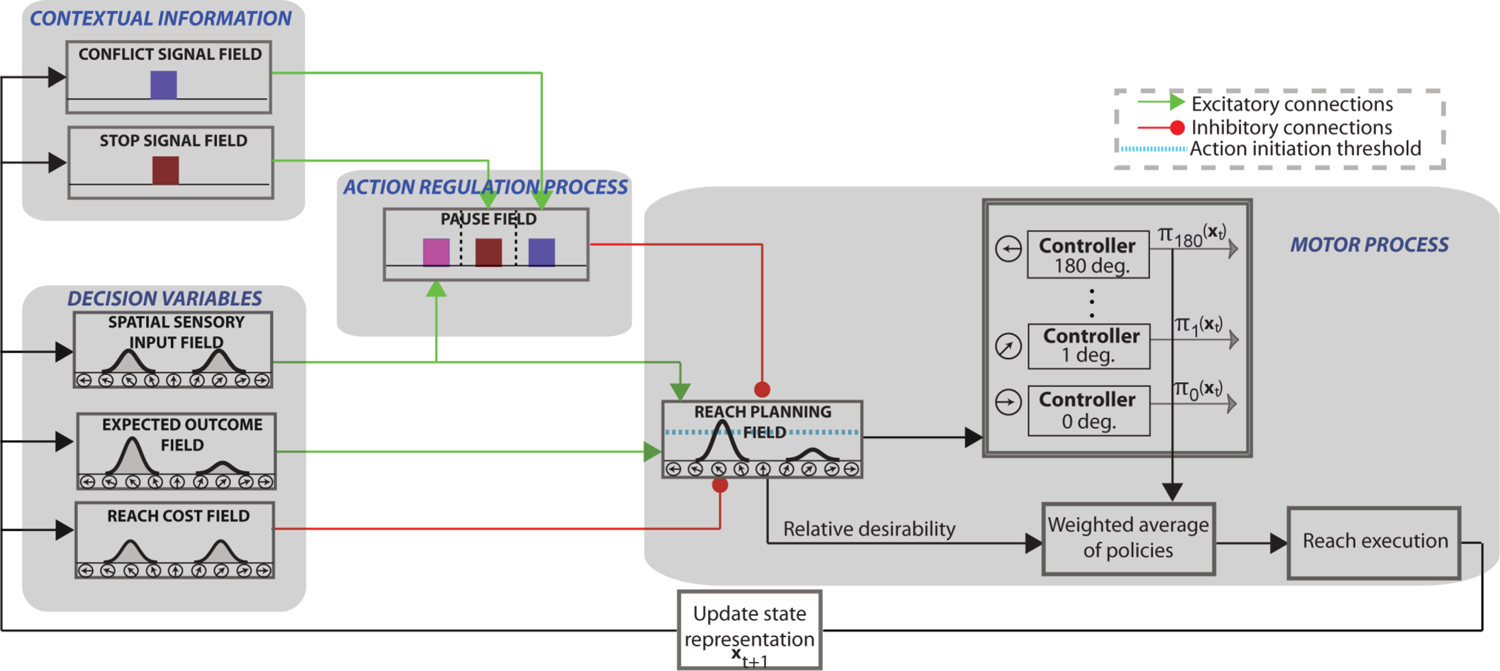
Model Architecture. The architectural organization of the neurodynamical theory to model tasks that involve action inhibition, such as decisions between competing options, decisions in the presence of conflicting information and outright stopping of actions.

We also added to the model architecture a Basal Ganglia (BG)-type mechanism for implementing the inhibitory process. This mechanism consists of three DNF platforms: (a) two context signal fields (stop and conflict) that represent information related to the contextual requirement of the tasks; (b) a pause field that suppresses the activity of the reach planning field to inhibit planned or ongoing actions. Each of the context fields consist of 100 neurons which project to the corresponding sub-population of the pause field via one-to-all excitatory connections. The stop signal field and the conflict signal field are activated when they detect a stop cue and conflict cue, respectively. Regarding the action selection function, the model does not need a context field to signal the decision task, since it can collect this information from the spatial sensory input field. In particular, the spatial sensory input field projects to the corresponding sub-population on the pause field with one-to-all excitatory connections. If more than one targets is encoded in the spatial sensory input field, the corresponding population on the pause field is triggered. Notably, this architecture is consistent with experimental studies which suggest dissociable frontal-BG circuits for different action suppression functions [34]. The pause field consists of 3 sub-populations of 75 neurons, each of them associated with one of the action regulation functions (i.e., action decisions between multiple options, action selection in the presence of conflicting information and outright stopping of actions). Once the pause field is triggered, the activity of the reach planning field is suppressed to delay a decision when more time is needed (i.e. during action selection or decision with conflicting information), or to completely suppress an action when it is no longer wanted or rendered inappropriate (i.e., outright stopping).

Each neuron in the reach planning field is connected with a stochastic optimal controller. Once the activity of a reach neuron *j* exceeds the action initiation threshold (cyan discontinuous line in Fig.4) at the current time and state ***x****_t_*, the corresponding controller initiates an optimal policy ***π****_j_*(***x****_t_*) to move the joystick towards the preferred direction of that neuron (see materials and methods section for more details). Reaching movements are generated as a mixture of active policies (i.e., policies in which the associated neuronal activity in the reach planning field is above the action initiation threshold) weighted by the normalized activity of the corresponding reaching neurons. The normalized activity is called relative desirability since it reflects the *attractiveness* of a policy with respect to alternatives (for more details see [19, 38].

#### 2.3.1 Modeling the computations of motor decision-making

The first task to model is the motor decision-making task that involves reaching to either a single direction (instructed trial) or selecting between two opposite directions (choice trial). Fig.5A illustrates the activity of the reach planning field as a function of time for a representative instructed (top panel) and choice (bottom panel) trial. Initially, the field activity is in the resting state. After the target onset in the choice trial, two neuronal populations selective for the targets are formed and compete through mutual inhibitory interactions. The activity of the pause field also increased to further inhibit the reach planning field to delay the initiation of the action (Fig. 6A blue trace shows the mean activity of the pause field across time in a choice trial). Once the activity of a neuronal population exceeds an action initiation threshold, the corresponding target is selected, the activity of the non-selected target is inhibited by the “winning” population, and a reaching movement is initiated. When only one target is presented (Fig.5A top panel), the activity of the corresponding neuronal population exceeds the action initiation threshold faster due to the lack of inhibitory competition from an alternative option and the non-activation of the pause field (Fig. 6A cyan trace shows that pause field activity remains on baseline). To get better insight on the model computations, consider two neurons in the choice trial, one from each population, centered at the target locations (Fig.5D). The neuron that exceeds the action initiation threshold first (red continuous traces) dictates the reaction time and the selected target (i.e., the selected direction of movement). In the absence of action competition (instructed trial), the activity of the reach neuron (blue trace) exceeds the action initiation threshold faster than when two actions compete for selection (red traces). Hence, we predict that simulated instructed reaches have shorter RT than reaches in the choice trials. To test this prediction, we simulated 100 decision-making trials in which 50 % of them involves choices between two competing options and the rest of them were instructed trials. Consistent with the prediction, we found that free choice movements have longer RT than instructed movements, as is shown in Fig.7A.

**Fig. 5.**
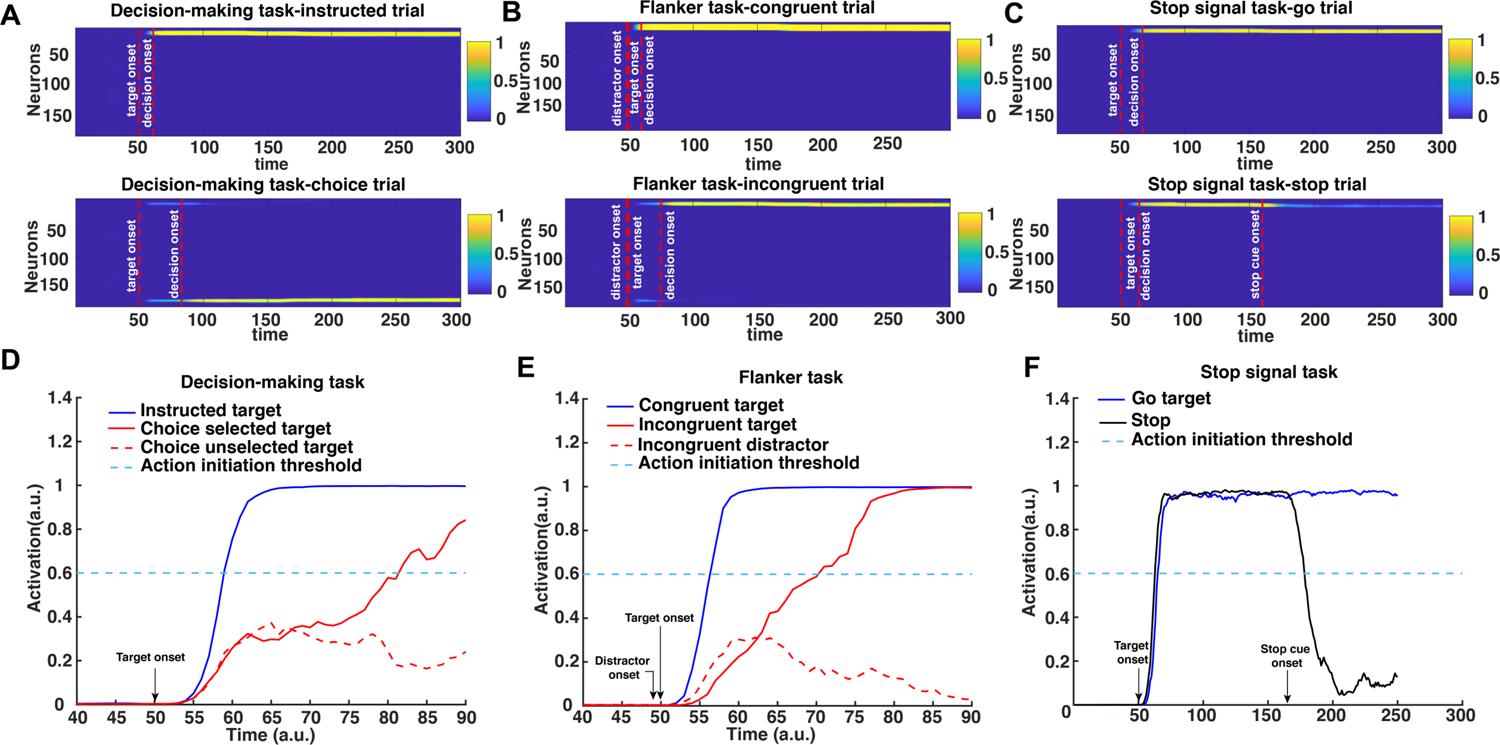
Simulated reach planning field neuronal activity changes in the decision making task, Eriksen flanker task and stop-signal task. (A)-(C) Activity changes of the 181 neurons in the reach planning field during the decision making task (instructed trial and choice trial)(A), the Eriksen flanker task (incongruent trial and congruent trial)(B), and the stop-signal task (go trial and stop trial)(C).(D)-(F) Activity changes of single neurons in the reach planning field during the decision making task(D), the Eriksen flanker task(E), and the stop-signal task(F).

**Fig. 6.**
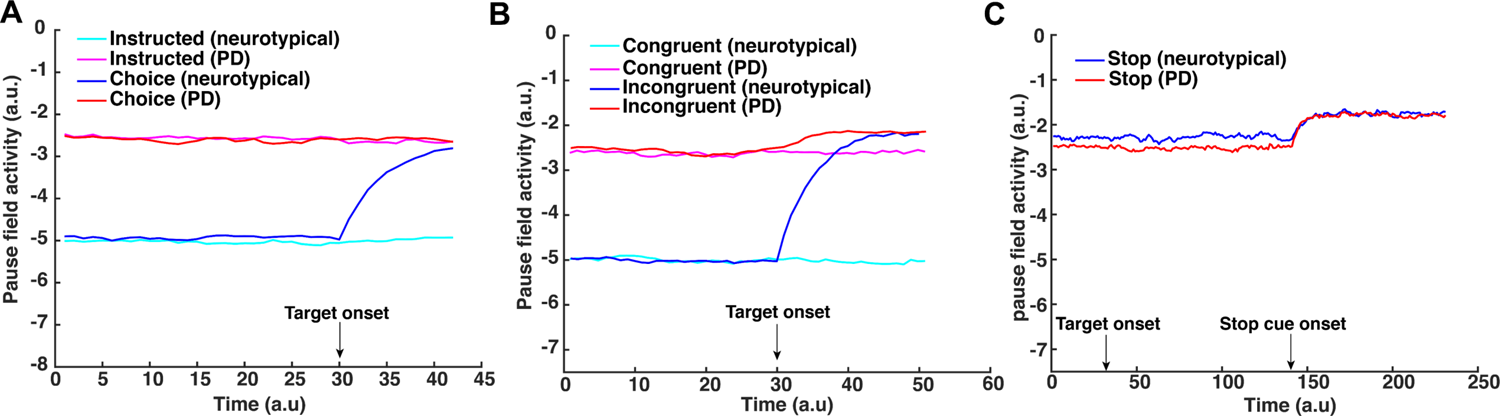
Simulated behavioral results from the three tasks. Simulated reaction time (RT) for the three experimental tasks predicted by the neurodynamical theory for both neurotypical individuals (Neurotypical, blue) and PD patients (PD, red).

**Fig. 7.**
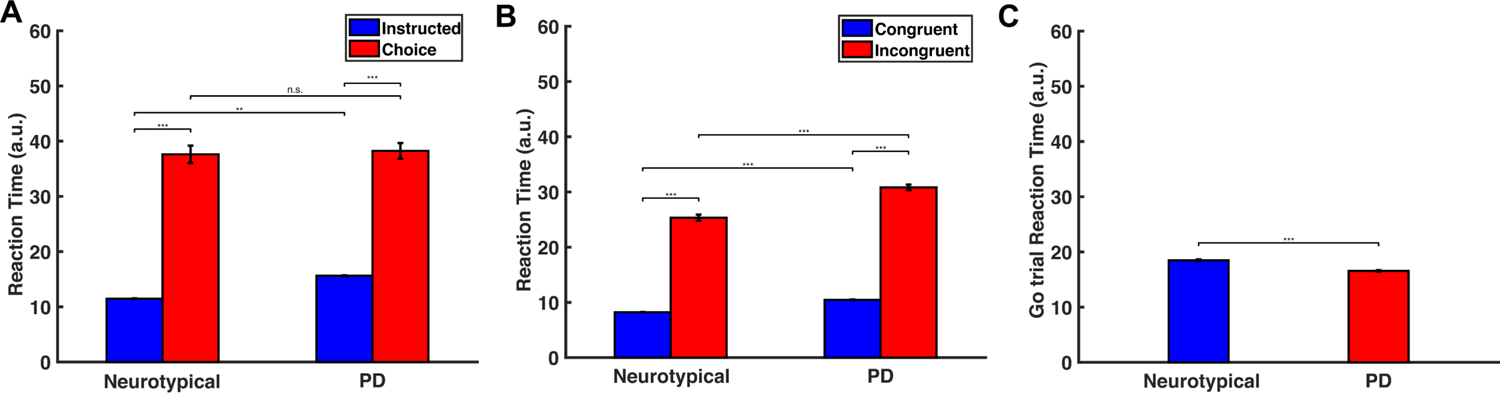
Simulated pause field activity changes during the three tasks. (A) Activity changes of single neuron in the pause field during the decision making-task. Cyan trace, simulated pause field activity during an instructed trial for a neurotypical individual. Magenta trace, simulated pause field activity during an instructed trial for a PD patient. Blue trace, simulated pause field activity during a choice trial for a neurotypical individual. Red trace, simulated pause field activity during a choice trial for a PD patient. (B) Activity changes of single neuron in the pause field during the Eriksen flanker task. Cyan trace, simulated pause field activity during a congruent trial for a neurotypical individual. Magenta trace, simulated pause field activity during a congruent trial for a PD patient. Blue trace, simulated pause field activity during an incongruent trial for a neurotypical individual. Red trace, simulated pause field activity during an incongruent trial for a PD patient. (C) Activity changes of single neuron in the pause field during the stop-signal task. Blue trace, simulated pause field activity during a stop trial for a neurotypical individual. Red trace, simulated pause field activity during a stop trial for a PD patient.

#### 2.3.2 Modeling the computations of conflicting information in motor decisions

In the Eriksen flanker task, a “flanker” (i.e., distractor) appears 100 ms before the target. Once the flanker is presented and detected by the spatial sensory input field, a reach neuronal population tuned to the flanker direction is formed - i.e., the model prepares an action towards the direction of the flanker. If the upcoming target coincides with the flanker direction (congruent trial), the pause field will not be activated (Fig.6B cyan trace) and the activity of the reach neuronal population will be further increased, leading to fast reaching movements towards the target direction (Fig.5B top panel). On the other hand, if the target points to the opposite direction from the flanker (incongruent trial), a new reach neuronal population is formed and competes with the reach neuronal population of the flanker (Fig.5B bottom panel). The conflict signal field detects the “conflicting information” and activates the pause field (Fig.6B blue trace) to suppress the reach planning field so that the target population will have time to further increase its activity and win the competition. The expected outcome field, which encodes the correct movement direction, biases the competition towards the target direction. To better understand the mechanism of action regulation in the Eriksen flanker task, we consider two neurons centered at the location of the target and the distractor, respectiely (Fig.5E). The neuronal activity of the distractor (red discontinuous trace) increases before the neuronal activity of the target (red continuous trace), since distractor precedes target presentation. Once the target is cued, the two neurons compete through inhibitory interactions. This competition, as well as the inhibition of the reaching neuronal population from the pause field, delay the action initiation, leading to longer RT. On the other hand, the lack of action competition and the non-activation of the pause field in the congruent trials (Fig.6B cyan trace) lead to shorter RT. To test this prediction, we simulated 100 Eriksen flanker task trials with 50 % of them to be incongruent trials. Consistent with the prediction, we found that reaching movements in incongruent trials have longer RT than in congruent trials as illustrated in Fig. 7B.

#### 2.3.3 Modeling the computations of outright stopping of actions

Regarding the stop-signal task, the model needs to generate actions while anticipating a stop signal. The experimental results showed that when people anticipate a stop signal, they have longer RT as compared to when they do not anticipate a stop signal (i.e., instructed trials). This suggests that the pause field is active even in the go trials to increase the chances of being able to abandon an action in case stopping is required. Τhe reach planning field activity in the go task resembles that of an instructed trial in the decision-making task (Fig.5C top panel), the only difference being that in the go trials the pause field is continuously active (Fig.6C blue trace). Hence, the activity of the reach planning field increases slower compared to the instructed trial, resulting in longer RT. In a go trial, the reach neuronal population tuned to the target direction is formed preparing an action. Once the activity of the population exceeds an action initiation threshold, the action is performed. However, in some trials, a stop signal is cued and the pause field activity is further increased, which subsequently inhibits the activity of the reach planning field to completely stop the planned or the ongoing action (Fig.5C bottom panel). To better understand the mechanism for stopping actions, consider one neuron from the population centered at the location of the target. The activity of the neuron increases once the target is cued and an action is initiated when the activity exceeds the action initiation threshold (Fig.5F blue trace). However, if a stop signal is cued, the pause field inhibits the activity of the neuron to stop the ongoing action (Fig.5F black trace). The stop signal is given with some delay (stop signal delay, SSD) in each trial. The longer the SSD is, the harder it is for the pause field to suppress the activity of the reach neuron increasing the chance to fail to stop the action. To test the model prediction, we simulated 50 go trials, as well as 250 stop trials, in which a stop stimulus appeared at different SSDs, signaling the model to abandon the action. Consistent with the model predictions, we found that go trials have longer RTs than instructed trials (p<0.001, two-way ANOVA, comparison made between the mean RT on instructed trials in the decision-making task and mean RT on go trials in the stop-signal task), and the probability to successfully stop a response reduces with increased SSD - i.e., the longer the signal delay, the harder it is for the model to stop an action (Fig.8 blue trace).

**Fig. 8.**
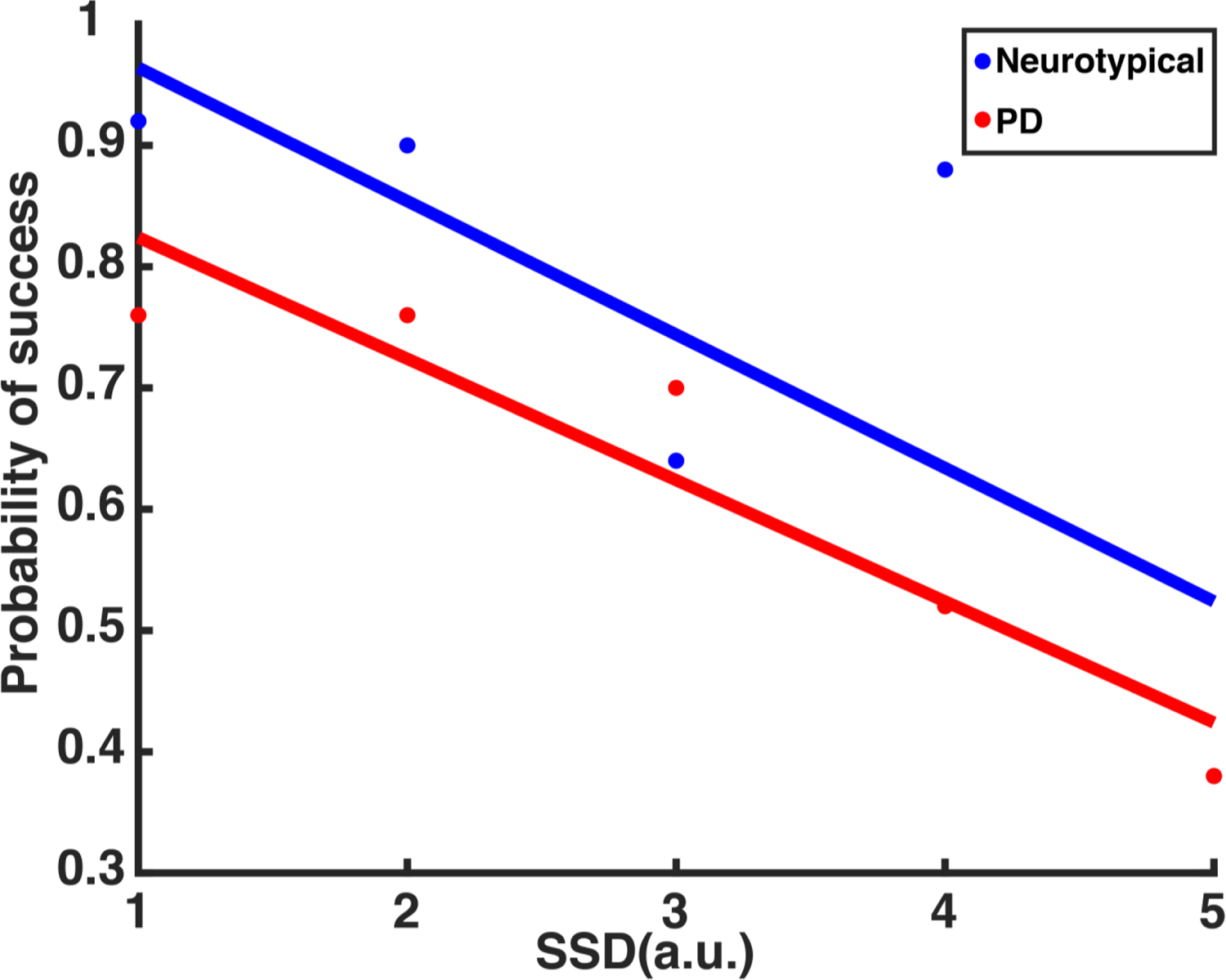
Simulated probability to successfully stop an action as a function of the stop signal delay (SSD) The (simulated) probability to successfully stop an action as a function of the SSD for neurotypical individuals (Neurotypical, blue) and PD patients (PD, red).

### 2.4 Dysfunction of the pause mechanism predicts motor impairment in PD patients

So far the neurodynamical theory is capable of capturing the motor behavior of the neurotypical participants in the 3 action regulation tasks. However, one of the main findings in our study is that PD patients exhibit overall slower responses in nearly all tasks compared to neurotypical participants. This motor impairment can be explained within the neurodynamical theory as a deficit on the pause mechanism. That is, the pause field is active even in the absence of conflicting information (congruent trials in the Eriksen flanker task) or competition between multiple actions (instructed trials in the decision-making task). To get a better understanding on how dysfunction of the pause mechanism affects motor behavior, Fig.6 shows the activity of the pause field as a function of time for a single trial across all tasks when simulating actions to model the RT of neurotypical participants and PD patients. In the decision-making task, the pause field is activated even when no action competition is presented (i.e., instructed trials) to capture the RT of the PD patient (Fig.6A magenta trace). This explains the slower response on initiating an action on instructed trials from PD patients. Also, the lack of difference on RT between neurotypical and PD patients in free choice trials suggests that the pause field exhibits similar activation levels when deciding between competing options after targets onset in both groups (Fig.6A). Regarding the flanker task, the pause field is active before the target onset in PD patients, explaining the slower response in both congruent and incongruent trials compared to neurotypical individuals (Fig.6B red and magenta traces). We need to point out here that although the pause field exhibits the same activation level in instructed and free choice trials (decision-making task) in PD patients, the slower response in choice trials compared to instructed trials is due to the inhibitory action competition between the two alternative movement directions.

Additionally, another important finding in our study is that PD patients have shorter RT in go trials than neurotypical participants in the stop signal task. By comparing the RT of movements between go trials in the stop-signal task and instructed trials in the decision-making task, we found that neurotypical participants delayed their responses in the go trials because they anticipated a stop signal as compared to when they did not anticipate a stop signal (i.e., instructed trials). This *response delay effect* (RDE) has been reported in previous studies [47–50] and has been associated with an “active braking mechanism” that increases the chance of abandoning a response in case stopping is required [51]. Note that PD patients also exhibited this active braking mechanism, but the difference in RT between go trials with anticipation of stopping signal and instructed trials was much smaller compared to neurotypical participants. Overall, these findings suggest that the pause field is active in the go trials for predicting the motor behavior in both groups of participants. In fact, the pause field activity is higher in neuropytical participants than PD patients, before a stop signal is detected, to explain the slower response of the first group compared to the latter group (Fig.6C, blue and red traces). We simulated 50 go trials with elevated pause field activity for both neurotypical participants and PD patients. Consistent with the behavioral findings, the go trials have longer RT in the simulated neurotypical participants than PD patients (Fig. 7C) (p<0.001, two sample t-test). Additionally, the model predicts that the probability to successfully stop a response is lower in PD patients than in neurotypical individuals (Fig. 8) due to the faster response of PD patients.

## 3 Discussion

### 3.1 General

Survival of species in an ever-changing environment requires flexibility in action selection. Traditional theories suggest that action selection takes place before action preparation [52–55]. However, recent cognitive theories challenge this view suggesting that in situations affording more than one alternative options, individuals prepare multiple actions in parallel that compete for selection before choosing one to execute [6–10]. This theory received empirical support from neurophysiological investigations in the sensorimotor areas of non-human primates (NHPs) showing that the brain encodes parallel reach, grasp and saccade plans before the animals select between them [11,12,56]. It is consistent with the continuous flow model of perception, which suggests that response preparation can begin even before the goal is fully identified and a decision is made [14–16]. Psychophysical support for this theory comes from the observation that when reaching to multiple potential targets, the initial movement is directed towards the average location of the targets, consistent with the theory that multiple prepared reaches being executed simultaneously [17–19].

Flexibility in action selection includes not only being fast and accurate enough when selecting between competing options, but also being flexible enough to change actions according to updated demands of the environment. This includes delaying actions in the presence of conflicting information and completely abandoning obsolete actions when they are rendered inappropriate [57–60]. Series of studies have explored how different brain regions contribute to programming, re-programming and stopping of actions using neural recordings and functional neuroimaging techniques [20, 21, 24, 25, 28, 29, 61, 62]. The basal ganglia (BG), and in particular, the subthalamic nucleus (STN), has been functionally implicated in action regulation, but in association with distinct frontal areas, such as the primary motor cortex (M1), the premotor cortex (preMC), the presupplementary motor area (preSMA) and the right inferior frontal gyrus (rIFG) [31–34]. In a sense STN seems to act as a “brake” when a stop signal is presented to rapidly suppress ongoing actions. Furthermore, various computational theories including the drift-diffusion model (DDM), urgency-gating model (UGM), evidence accumulation models (EAMs), race models and mutual inhibition models, have been constructed to explain how the brain selects between competing options, inhibits actions in the presence of conflicting information and abandons planned or ongoing action when they are rendered inappropriate [63–66]. Although these theories provide significant insights into the action regulation mechanisms, a major limitation is that they explored separately each of these three motor functions, making it challenging to develop a unified theory of action regulation. A computational theory that can simulate the mechanisms underlying selecting, inhibiting and outright stopping of actions is needed to unify and integrate these distinctly studied actions and mechanisms. Our research focuses exactly on what has been missing from previous studies – to design a large scale computational theory that can predict: 1) how the brain selects between competing actions, delays actions in the presence of conflicting information and stops actions when they are rendered inappropriate, 2) how neuropsychiatric diseases, such as PD, affect the action regulation circuitry and lead to motor deficits. Building on our previous work [38, 39], we developed a neurodynamical framework to integrate the three action regulation functions into a unified computational theory. This computational theory is based on the affordance competition hypothesis, in which multiple actions are formed concurrently and compete over time until one has sufficient evidence to win the competition [6]. We replace evidence accumulation with desirability – a continuously evolving quantity that integrates all sources of information about the relative value of an action with respect to alternatives. The winning action determines the reaction time and the direction of movement. The computational theory includes a BG-type mechanism of inhibiting actions in the presence of competing options, conflicting information and stopping signals. We tested the computational model in a series of tasks that involve action selection, decision conflict and outright stopping using neurotypical individuals and PD patients. Our findings showed that the model captures many aspects on human behavior, such as the longer RT in the presence of competing actions and conflicting information, as well as the inverse relationship between the probability to successfully stop an action and stop signal delay (SSD). It also predicts the motor impairment on PD patients when performing these three motor tasks as a deficit in the pause mechanism. In particular, the model explains the longer responses in generating actions even without the presence of competing action and conflicting information in PD patients compared to neurotypical participants as a consequence of hyperactivity on the pause field. This is consistent with experimental evidence showing that STN is overacting in PD patients [67] leading to longer responses in visuomotor tasks. Overall, to the best of our knowledge, our study presents the first neuro-computational theory that integrates the mechanisms of three action regulation functions and predicts how disruption of these mechanisms can lead to motor deficits reported in neurological diseases such as PD.

### 3.2 Mapping to neurophysiology

The computational theory presented in the current study is a systems-level framework aimed to qualitatively predict response patterns of neuronal activities in ensembles of neurons, as well as motor behavior, in action regulation tasks. It captures many key features of the functional properties of the cortical-subcortical network involved in action regulation. The spatial sensory input field mimics the organization level of the posterior parietal cortex (PPC) [68, 69]. The expected reward field can be associated with the ventromedial prefrontal cortex (vmPFC) and orbitofrontal cortex (OFC), two frontal areas with important role in computation and integration of reward [70, 71]. The reach cost field can be equated to the anterior cingulate cortex (ACC) that has a key role in computing the cost for performing an action [72, 73]. The reach planning field can be associated with the parietal reach region (PRR) [74, 75] and the premotor dorsal cortex (PMd) [76, 77], two cortical areas involved in planning of reaching movements. The stop signal field can be equated with the right inferior frontal gyrus (rIFG), which is recruited when cues associated with response inhibition are detected [78, 79]. Regarding the conflict signal field, the popular view is that the pre-supplementary motor area (preSMA) detects the co-activation of different but conflicted responses (e.g., naming the color of the word *red* written with green color), it activates the STN to *temporarily* suppress a response [80, 81]. Finally the pause field can be equated to the STN which is activated in tasks that require stopping or pausing behavioral outputs to suppress actions [23, 27, 30, 35].

### 3.3 Computational modeling of action inhibition deficits in PD patients

PD is a progressive neurodegenerative disease associated with progressive loss of dopaminergic neurons in the substantia nigra of the BG [67]. The disruption of frontal-BG circuitry is responsible for the development of major symptoms of PD, including rigidity, tremor, bradykinesia, and postural instability [82, 83]. In particular, impairment of response inhibition abilities, which greatly affects the life quality of PD patients, is considered to be a sensitive measure to the progression of PD [84]. As a key player in the frontal-BG circuit, the STN is suggested to mediate a “pause” function by rapidly inhibiting the BG activity. Therefore, it is considered to play a prominent role in the pathology of PD [85]. An increase in the neuronal activity of the STN has been demonstrated in electrophysiological and behavioral studies in non-human primate models of PD [86] as well as PD patients [87]. Our findings suggest that an increase in the STN-mediated “pause” signal is responsible for the impairment of action inhibition abilities in PD patients. In our model, we assigned higher baseline activation level of the pause field in the decision-making task and Eriksen flanker task in PD patients compared with neurotypical participants. Consistent with the model predictions, PD patients exhibited longer RT than healthy individuals in the instructed trials of the decision making task, as well as in both incongruent trials and congruent trials of the Eriksen flanker task. Notably, RT in the free choice trials of the decision-making task wasn’t significantly different between PD patients and neurotypical participants. This suggests that pause field, which is already highly active in instructed trials, is not further activated in the choice trials.

An interesting finding in our study was that neurotypical individuals had slower response than PD patients to initiate an action in the stop-signal task. This is somewhat counter-intuitive since the STN is overactive in PD patients and therefore we would expect that they would had slower response than neurotypical participants. However, when we compared RT between instructed trials (decision-making task) and go trials (stop signal task) of the neurotypical individuals, we found that they responded slower when they anticipated a stop signal. On the other hand, we found much subtler difference in RT between instructed trials and go trials in PD patients. This suggests that the pause mechanism is activated in the stop-signal task in neurotypical individuals even before a stop signal is presented. By activating the pause field to simulate the motor behavior of the neurotypical participants, the model predicts that PD patients will have faster responses and lower probability to stop planned or ongoing actions compared to neurotypical participants. In other words, the model explains the slower responses of the neurotypical participants as a cognitive strategy adapted to minimize the probability to fail to stop an action if a stop signal is detected.

### 3.4 Activity suppression or increase of action initiation threshold?

In our theory, the pause field delays motor decisions by suppressing the activity of the reach planning field. However, an alternative hypothesis is that the pause field mediates the action inhibition function by increasing the action initiation threshold. Previous studies suggest that STN low-frequency oscillatory activity and medial prefrontal cortex (mPFC)-STN coupling are involved in determining the amount of evidence (i.e., action initiation threshold) needed before making a decision [88–91]. Additionally, clinical studies showed that deep brain stimulation targeting the STN in PD patients can modulate the amount of evidence, and therefore the action initiation threshold, required to initiate an action [89]. Hence, it is also likely that the STN delays motor decisions in the presence of competing actions and/or conflicting information by increasing the action initiation threshold, instead of suppressing the activity of the motor areas that generate actions. Our computational theory is capable of modeling this hypothesis by adjusting the action initiation threshold in the reach planning field. However, it cannot dissociate between the two hypotheses on how the STN pauses actions when needed. To do so, future neurophysiological or neuroimaging studies need to record activity from the STN and motor areas during decision tasks with multiple options and/or conflicting information.

### 3.5 Hyperactive pause mechanism or altered cost/reward ratio in PD patients?

Although our study suggests that deficits in movement preparation in PD patients, such as slow reaction times, are related to hyperactivity in STN that inhibits planned actions, other studies have associated motor impairments with motivational deficits [92]. In particular, motivational deficits seem to significantly contribute to bradykinesia in PD patients and lead to alternation in the amount of effort required to perform a movement at normal speed, as well as the perceived reward for successfully completing the action [93].

Motor decisions are frequently made based on expected reward and the associated effort cost required to obtain the reward. The cost has been considered to be detrimental, since we tend to choose the less costly actions especially when they are associated with similar expected rewards [94, 95]. The dopaminergic neurons seem to be critically involved in the process of cost versus reward (i.e., cost/reward ratio) evaluation. Dopamine depletion from rat results in decreased tolerance for effort cost, whereas enhanced dopamine levels has the opposite effect [94, 96]. Loss of dopaminergic neurons and their projections is a major pathological hallmark in PD patients. Clinical studies have shown that PD patients, regardless of medication status, tend to engage less effort for the lowest reward compared with neurotypical participants in a hand-squeezing task [93]. However, dopamine medication motivates PD patients to engage more effort for a given reward, comparing to their off medication state. In addition, Deep brain stimulation (DBS) of the STN establishes a reliable congruency between action and reward in PD patients and remarkably enhances it over the level observed in neurotypical individuals [97].

Overall, these studies provide evidence that impairment of movement preparation in PD patients can also be related to deficits in the mechanism that evaluates reward and effort cost associated with actions - i.e., alternation of the cost/reward ratio. Notably, this can be also modeled within our neurodynamical framework by increasing the amount of effort required to perform actions towards the cued directions. Additionally, the alternation of the cost/reward ratio in PD patients could be also related to the hyperactivity of the STN - more effort is required to increase the activity of the motor population, which is continuously inhibited by the STN, to initiate an action. Today, the mechanism for motor and information processing deficits in PD patients is still under extensive study. PD is considered not only a disease caused by degeneration of substantia nigra dopaminergic neurons, but also a system-level disease caused by dysfunction of the cortical-BG circuit [67]. Therefore, both the hyperactivity of the STN and the altered cost/reward ratio can be considered parts of PD pathophysiology, and contribute to the motor deficits in PD patients.

### 3.6 Conclusions

In conclusion, the current study aims to advance our understanding on the computations underlying action regulations in tasks that involve action inhibition, the failure of which contributes to various neuropsychiatric diseases. We proposed a large scale neurodynamical computational framework that combines dynamic neural field theory with stochastic optimal control theory to simulate the mechanisms of action regulation and to predict how disruption of this mechanism lead to motor deficits in PD patients. We evaluated the model predictions by comparing the motor behavior of neurotypical individuals and PD patients in three tasks that require action inhibition. To the best of our knowledge, our results revealed for the first time an integrated mechanism of action regulation that affects both action planning and action inhibition. When this mechanism is disrupted (as in PD patients), motor behavior is affected, leading to longer reaction times and higher error rates in decisions and actions. Overall, our findings provide significant insight on how the brain regulates actions that involve inhibition, and open new avenues for improving and developing therapeutic interventions for diseases that may involve these circuits.

## 4 Methods

### 4.1 Participants

A sample of 15 adults with PD and 32 neurologically healthy, age-matched adults took part in the study. The study was approved by the University of California, Los Angeles Review Board and all individuals signed an informed consent before participating.

### 4.2 Stimuli and Procedure

#### 4.2.1 Decision-Making Task

All experiments were programmed using Psychtoolbox 3 for Matlab. Experimental setup is shown in Figure 1. In the decision-making task, participants sat in a dark room in front of a 22-inch Dell LED monitor where stimuli would be presented on. The screen was approximately 50 cm away from the participant. A two-dimensional joystick (Thrustmaster T.16000M FCS, maximum range of axis value is −32,000 +32,000) was placed in front of the monitor. During the task, the participants were instructed to move the joystick towards the left or right direction using their right hand in reaction to the corresponding stimulus. Each trial started with the screen turning black. After 1.0-1.1 s, a white fixation cross appeared in the center of the black screen for 1.0-1.1 s, then the white fixation cross disappeared, and four white arrows appear in the center of the black screen. In 50% of the trials (choice trials), two of the arrows pointed to the left, and the other two to the right (e.g. < < > >), in which case the participant needed to freely decide whether they would move the joystick to the left or right. In the other 50% of the trials (instructed trials), the four arrows were pointing to the same direction (left or right) (e.g. < < < <), in which case the participant needed to move the joystick towards the direction the arrows were pointing to. The arrows remained on the screen for up to 1.5 s before they disappear, then the screen turned black for 0.5 s. If the participant responded to the stimulus by moving the joystick to the left or right (axis value threshold for response: −25000 to the left/+25000 to the right) when the arrows were presented on the screen, after 10ms, the screen would turn black for the remaining of the 1.5 s plus 0.5 s, after which the screen would remain black and the next trial would start. If the joystick did not return to the baseline (axis value between −2500 and +2500), the next trial would not start until the joystick returned to the baseline. Every participant performed 2 blocks of trials, with 52 trials in each block. In each block of trials, there are 26 choice trials and 26 instructed trials. The trial type (choice or instructed) were randomized. Before each trial, the participant did not know whether the next trial would be a choice trial or an instructed trial.The RT for each trial was recorded as the time interval between the appearance of the arrows on the screen and the participant’s response.

#### 4.2.2 Eriksen flanker Task

An arrow version of the Eriksen flanker Task [98] with arrows pointing to the left and right was performed in our study. During the Eriksen Flanker task, the same equipment as described in 3.2.1 were used, the major difference being that in each trial, the target stimulus was flanked by stimuli which were pointing to the opposite direction of the target arrow (incongruent trial) or to the same direction as the target arrow (congruent trial), and every participant was told to move the joystick towards the same direction as the target arrow using his/her right hand. In each trial, the screen first turned black for 1.0-1.1 s, then a white fixation cross appeared in the center of the screen for 1.0-1.1 s. After this interval, four white flanker arrows pointing to one direction (left or right) appeared in the center of the screen, leaving a blank space in the middle (e.g. < < < <). After 100 ms, a white target arrow appeared in the blank space, pointing either to the opposite direction of the flankers (incongruent trial) or the same direction (congruent trial). The target arrow and the flankers remained on the screen for up to 1.5 s, then disappeared, and the screen turned black for 0.5 s. If the participant responded to the target arrow by moving the joystick to the left or right, after 10ms, the screen would turn black for the remaining of the 1.5 s plus 0.5 s, after which if the joystick returned to baseline, the screen would remain black and start the next trial. Each participant performed two blocks of trials, with 52 trials in each block, making a total of 104 trials. In each block of trials, there are 26 incongruent trials and 26 congruent trials. The direction of the target arrows and the type of flanker (incongruent or congruent) were randomized. The RT for each trial was recorded as the time interval between the appearance of the target arrow and the participant’s response.

#### 4.2.3 Stop Signal Task

A trial in a stop signal task is either a go trial or a stop trial. In each trial, arrows pointing to the left or right direction were presented on the screen as a stimulus. In a go trial (no stop signal is presented), the participant should respond as fast as possible by moving the joystick towards the direction the arrows were pointing to. In a stop trial, the participant should try to inhibit their response after the stop signal was cued. Participants were told that stop was not always possible, and that stop trials and go trials are equally important. Before the experiment, each participant performed 24 training trials, including 16 go trials and 8 stop trials. At the beginning of a trial, the screen turned black. After 1.0-1.1 s, a white fixation cross appeared in the center of the screen for 1.0-1.1 s, then the fixation cross disappeared, and four white arrows pointing to the left or right appeared in the center of the screen. In a go trial, the arrows remained on the screen for up to 1.5 s before they disappeared, then the screen turned black for 0.5 s. If the participant responded to the stimulus by moving the joystick when the arrows were presented on the screen, after 10ms, the screen turned black for the remaining of the 1.5 s plus 0.5 s, after which if the joystick returned to baseline, the screen remained black and the next trial was started. A stop trial is nearly identical to a go trial, except that the arrows turned red after an interval termed “stop signal delay” (SSD) indicating that the participant should abandon any response immediately. If the participant inhibited their actions, the arrows remained on the screen for the rest of 1.5 s, and in the subsequent stop trial, the SSD would increase by 50 ms, making inhibition more challenging. If the participant failed to inhibit their actions, after 10 ms, the arrows disappeared, and the screen turned black for the remaining of the 1.5 s plus 0.5 s, after which if the joystick returned to the baseline, the screen remained black and the next trial would start. In this case, the SSD would decrease by 50 ms, making it easier to inhibit actions. Each participant performed 3 blocks of trials, with 60 trials in each block. In each block of trials, there were 40 go trials and 20 stop trials. The direction of the arrows and the type of trial (go or stop) were randomized. The RT for each go trial and failed stop trial were recorded as the time interval between the appearance of white arrows and the participant’s response.

### 4.3 Statistical Analysis

Cubic interpolating splines were used to smooth the reach trajectories and compute the velocity of the movements. Reaction time (RT) was defined as the time between the target appearance and the time that reach velocity exceeded 10% of the maximum velocity. RTs faster than 100ms were removed because anticipation is considered to be involved prior to actions, as well as RTs longer than 1500ms. RT outliers (RTs >3 standard deviations below or above the mean RT) were also excluded from the analysis. The trials in which the participant changed their mind (moving towards one direction past 5% of the maximum range, and then changed their mind to move towards the other direction) were also excluded from further analysis. RTs across all participants were pooled together, and for the decision making task and the Eriksen flanker task, two-way ANOVA analysis was performed to determine the group differences in RTs. For the stop signal task, two-sample t-test was performed to determine the group differences in go trial RTs.

### 4.4 Computational Model Architecture

We developed a neurodynamical framework based on our previous studies [38, 39] to model the three action regulation functions. The computational framework combines dynamic neural field (DNF) theory with stochastic optimal control theory, and includes circuitry for perception, expected outcome, effort cost, context signal, pause, action planning and execution. Each DNF simulates the dynamic evolution of firing rate activity of a network of neurons over a continuous space with local excitation and surround inhibition. The functional properties of each DNF are determined by the lateral inhibition within the field and the connections with other fields in the architecture. The projections between the fields can be topologically organized – i.e., each neuron i in the field drives the activation at the corresponding neuron i in the other field (one-to-one connections), or unordered – i.e., each neuron in one field is connected with all neurons on the other field (one-to-all connections). The activity of a field *j* evolves over time under the influence of external inputs, local excitation and lateral inhibition interactions within the field, as well as interactions with other *k* fields, as described by Equation (1):

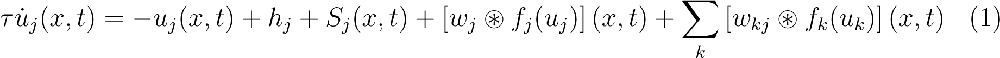

where *u*(*x, t*) is the local activity of the DNF at the position *x* and time *t*, and *ů_j_* (*x, t*) is the rate of change of the activity over time scale by a time constant *τ*. If there is no external input *S*(*x, t*), the field converges over time to the resting state *h* from the current level of activation. The first convolution term 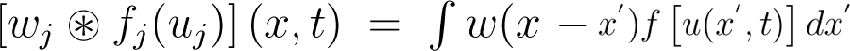 models interactions between the simulated neurons at different locations within the field j, and is shaped by the interaction kernel of Equation (2), which consists of both excitatory and inhibitory components:

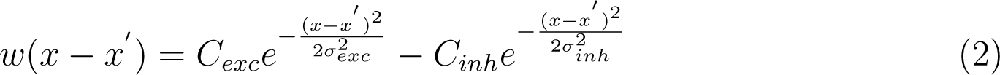

where *C_exc_*, *C_inh_*, *σ_exc_* and *σ_inh_* describe the amplitude and the width of the excitatory and the inhibitory components, respectively. We convolved the kernel function with a sigmoidal transformation of the field so that the neurons with activity above a threshold participate in the intra-field interactions:

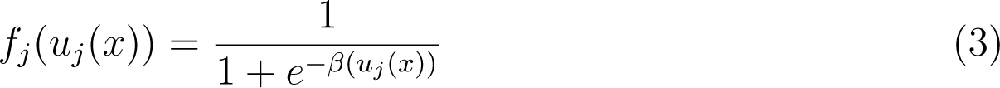

in which the steepness of the sigmoid function was controlled by *β*.

The function *w_jk_* describes the connectivity kernel between fields *u_j_* and *u_k_* showing the contribution of field *u_k_* to the dynamics of field *u_j_*. The sigmoid *f_k_*(*u_k_*) and *w_jk_* are convolved to determine the full contribution from field *u_k_* to *u_j_*.

The architectural organization of the framework is shown in Figure 4. The “reach planning” field encodes the potential movement directions, and is responsible for initiating the reaching movements. The “spatial sensory input” field encodes the angular representations of the competing targets. The “expected outcome” field encodes the expected reward for reaching to a particular direction centered on the hand position. The outputs of these two fields send excitatory projections (green arrows) to the reach planning field in a topological manner. The “reach cost” field encodes the effort cost required to implement an action at a given time and state. The reach cost field sends inhibitory projections (red arrow) to the reach planning field to penalize high-effort actions. For instance, an action that requires changing of moving direction is more “costly” than an action of keep going in the same direction. The “pause” field suppresses the activity of the reach planning field to inhibit planned or ongoing actions via inhibitory projections to the reach planning field. The stop signal field and the conflict signal field encode information related to the contextual requirement of the task (i.e., stopping cue or flanker distractor), and send one-to-all excitatory projections to the corresponding population of the pause field. In particular, the stop signal field is projected to the neuronal population of the pause field which is responsible for outright stopping of action, whereas the conflict signal field projects to the neuronal population of the pause field, which is responsible for delaying decisions when conflicting information is detected. Each of the context signal fields (stop signal field and conflict signal field) consists of 100 neurons, whereas the pause field consists of 3 neuronal sub-populations, each consists of 75 neurons. The rest of the fields consist of 181 neurons with a preferred direction between 0 to 180 degrees. The activity of the reach planning field *S_action_* is given as the sum of the outputs of the fields encoding the position of the target *v_pos_*, the expected reward *v_reward_*, the estimated reach cost *v_cost_*, and the activity from the pause field *v_pau_*, at any given time and state, corrupted by a Gaussian distributed additive noise *ξ*.

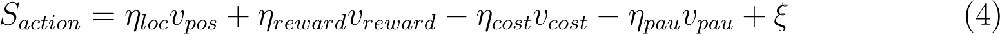

where *η_loc_*, *η_reward_*, *η_cost_* and *η_pau_* are scalar values that weigh the influence of the spatial sensory input field, the expected outcome field, the reach cost and the pause field, respectively, to the activity of the reach planning field. The values of the model parameters are given in S1 Table. The normalized activity of the reach planning field describes the relative desirability *d_i_* of each “reach neuron” with respect to the alternative options at time t – i.e., the higher the activity of a reach neuron *j*, the higher the desirability to move towards the preferred direction *φ_j_* of this neuron with respect to the alternatives at a given time t. Each neuron *j* in the reach planning field is connected with a control scheme that generates reaching trajectories. Once the activity of that neuron exceeds the action initiation threshold *γ*, the controller is triggered and generates an optimal policy ***π_j_***, a sequence of motor actions towards the preferred direction of the neuron *j*. The optimal policy is given by minimization of the cost function:

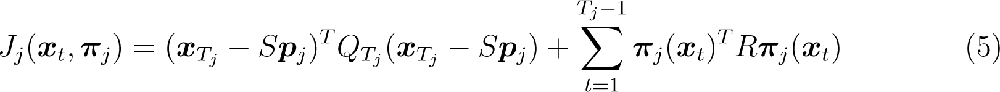

where ***π****_j_*(***x****_t_*) is the policy from the time t = 1 to t = *T_J_* to reach towards the preferred direction *φ_j_*; *T_j_* is the time required to arrive at position ***p_j_***; ***p_j_*** is the position planned to arrive (goal position) at the end of the reaching movement, given by ***p_j_*** = [rcos(*φ_j_*), rsin(*φ_j_*)], in which r is the distance between the current location of the hand and the location of the stimulus encoded by the neuron *j*. ***x****_Tj_* is the state vector at the end of the reaching movement, and matrix S selects the actual position of the hand and the goal position at the end of the reaching movement from the state vector. Matrices *Q_Tj_* and R define the cost dependent on precision and control, respectively. More details about the optimal control model are described in [38, 39]. Consequently, a action is initiated once a neuronal population exceeds the action initiation threshold and the executed action ***π****_mix_*(***x_t_***) is given as a mixture of the active policies (i.e., policies with active neurons) weighted by relative desirability values of the corresponding neurons at any given time and state.

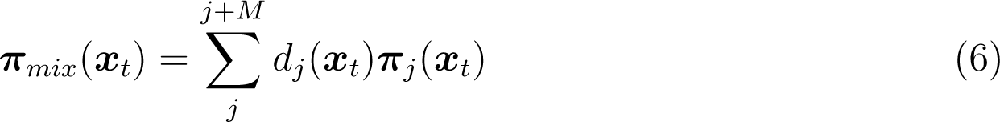

where ***x_t_*** is the state of the system at time *t* (i.e., position, velocity, orientation of the trajectory), *d_j_* is the normalized activity of the neuron *j* (i.e., relative desirability value of the neuron *j*), and ***π****_j_* is the optimal policy generated by the controller connected with neuron *j*. Because desirability is time- and state-dependent, the weighted mixture of the individual policies can change/correct the current trajectory in the presence of new incoming information - e.g., a stop signal cued while acting. In order to handle contingencies during the movement, the “receding horizon control”(RHC) [99, 100] technique, also known as model predictive control (MPC), which is widely used in stochastic optimal control models, was implemented in the framework. According to RHC, the framework would only execute the initial portion of the sequence of actions for a short period of time *τ* (*τ* = 9 in our framework), after which the framework would recompute the optimal policy ***π****_mix_*(***x****_t_* + *τ*) from time t+*τ* to t+*τ* +*T_i_*, and this approach would continue until the hand reaches one of the targets.

## Acknowledgments

Research reported in this publication was supported by National Institute of Neurological Disease and Stroke of the National Institutes of Health under award number U01NS098961 and R01NS097782. The content is solely the responsibility of the authors and does not necessarily represent the official views of the National Institutes of Health.

## Author Contributions

N.P. and V.N.C conceived the study.; J.C., M.M, N.P. and V.N.C designed the experiment; N.H., D.B. and M.M. recruited subjects and collected the data; S.Z. and J.C. performed the data analysis; S.Z designed the neurocomputational model and performed the simulations; S.Z drafted the manuscript with substantial contribution from N.P. and V.N.C; S.Z, J.C., N.P. and V.N.C. revised and approved the manuscript.

## Data Availability

All relevant data are within the manuscript and its supporting information files and will be uploaded to the Open Science Framework (OSF) upon approval for publication.

---

**S1 Table.**

| **Model Parameters** |  |  |
| --- | --- | --- |
| **Parameters** | **Description** | **Value** |
| $\eta_{loc}$ | Visual input gain | 8.5 |
| $\eta_{reward}$ | Expected outcome input gain | 2.5 |
| $\eta_{cost}$ | Action cost input gain | -0.1 |
| $\eta_{pau}$ | Pause input gain | -4.0 |
| $\gamma$ | Action initiation threshold | 0.6 |

| **Spatial sensory input field & Expected outcome field parameters** |  |  |
| --- | --- | --- |
| **Parameters** | **Description** | **Value** |
| $\tau$ | Time constant | 5.0 |
| $c_{exc}$ | Amplitude of excitatory portion of weight kernel | 0 |
| $c_{inh}$ | Amplitude of inhibitory portion of weight kernel | 0 |
| $\sigma_{exc}$ | Width of excitatory portion of weight kernel | 5.0 |
| $\sigma_{inh}$ | Width of inhibitory portion of weight kernel | 40.0 |
| $h$ | Resting activity level | -5.0 |
| $q$ | Noise level | 0.25 |
| $\sigma_q$ | Width of noise kernel | 5.0 |
| $\beta$ | Steepness of sigmoid activity function | 1.0 |

| **Reach planning field parameters** |  |  |
| --- | --- | --- |
| **Parameters** | **Description** | **Value** |
| $\tau$ | Time constant | 5.0 |
| $c_{exc}$ | Amplitude of excitatory portion of weight kernel | 0 |
| $c_{inh}$ | Amplitude of inhibitory portion of weight kernel | 20 |
| $\sigma_{exc}$ | Width of excitatory portion of weight kernel | 5.0 |
| $\sigma_{inh}$ | Width of inhibitory portion of weight kernel | 180 |
| $h$ | Resting activity level | -5.0 |
| $q$ | Noise level | 0.5 |
| $\sigma_q$ | Width of noise kernel | 5.0 |
| $\beta$ | Steepness of sigmoid activity function | 1.0 |

| Pause field parameters |  |  |
| --- | --- | --- |
| Parameters | Description | Value |
| $\tau$ | Time constant | 5.0 |
| $c_{exc}$ | Amplitude of excitatory portion of weight kernel | 0 |
| $c_{inh}$ | Amplitude of inhibitory portion of weight kernel | 0 |
| $\sigma_{exc}$ | Width of excitatory portion of weight kernel | 5.0 |
| $\sigma_{inh}$ | Width of inhibitory portion of weight kernel | 25.0 |
| $h$ | Resting activity level | -5.0 |
| $q$ | Noise level | 0.25 |
| $\sigma_q$ | Width of noise kernel | 5.0 |
| $\beta$ | Steepness of sigmoid activity function | 1.0 |

